# Helping those who help others for indirect fitness benefits not indirect reciprocity

**DOI:** 10.1101/2019.12.11.872937

**Authors:** Gilbert Roberts

## Abstract

Helping those who help others appears to be a widespread phenomenon. It is typically framed as indirect reciprocity in which individuals who are seen to help later receive returns from third parties. However, indirect reciprocity only works when individuals condition their help not just on how their recipient has behaved in the past but also on whether their recipient was justified in behaving that way. It also requires sufficient repeated interactions of this type among other individuals for a benefit to be reciprocated. These factors limit the scope of indirect reciprocity to explain cases where people do help those who help others. Here, I propose instead that helping can be explained by the indirect fitness benefits (or ‘relatedness’) that result from helping other helpers in groups. This means that when individuals help other helpers, they may not make any returns via indirect reciprocity, but rather they may be helping a strategy of helping those who help. In this way, the helping strategy can spread even when helping has no net benefit to the individual helper. This is a form of relatedness in which individuals help their kin that are recognized by their helping behaviour. As such, conditional helping is likely to be found where population structure promotes relatedness through non-random association. The analysis suggests indirect reciprocity may not have played the decisive role in the evolution of human cooperation that is often thought, but paradoxically that the use of image scores deserves renewed attention as a strategy of helping those with the same behaviour.

## Introduction

Explaining how cooperation works outside of the confines of directly reciprocal relationships and close familial ties, especially in large-scale human societies, has become a major focus of research. A popular conception is that we tend to help those who help others (Nowak & Sigmund, 2005). There is some evidence for this and it is interpreted in terms of indirect reciprocity (IR) (Wedekind & Milinski, 2000). Here, I point out that the focus on IR is misplaced and that helping those who help others is selected for through indirect fitness benefits (Hamilton, 1964).

IR appears to map onto the notion of helping those who help others because it involves one individual paying a cost to benefit another, and then a third party paying a cost to benefit the original donor (Boyd & Richerson, 1989). Provided the benefits exceed the costs then both make a net profit (Figure 1). IR is sometimes wrongly thought of as an umbrella term for fitness benefits arising through third parties (Roberts et al., 2021) so it is important to remember that it is a specific mechanism that depends on repeated interactions among third parties (Kandori, 1992). The best known advance in the theory of IR was the concept of ‘image scoring’ (Nowak & Sigmund, 1998): individuals who help others acquire a good image score and potential donors use these image scores to decide whether to help. Simulations showed that this could lead to mutually beneficial cooperation without donors ever meeting their recipients again. This appears to be supported by experimental evidence that people give preferentially to those with positive image scores (Milinski, 2016; Milinski et al., 2001; Seinen & Schram, 2006; Semmann et al., 2004; Wedekind & Milinski, 2000).

**Figure 1.**
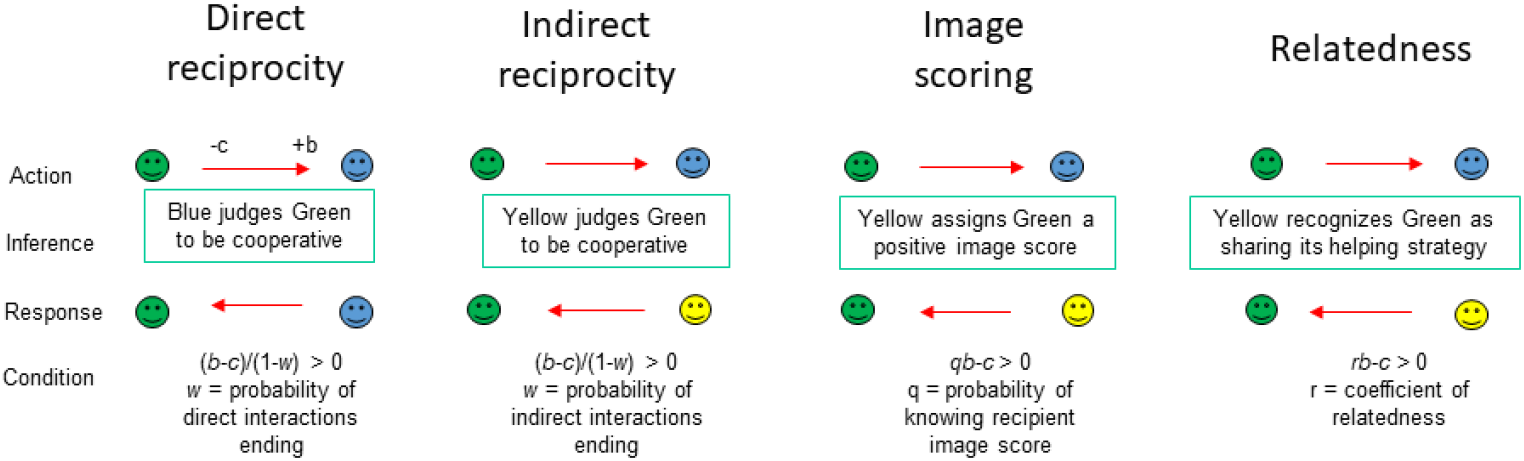
Reciprocity and relatedness. A comparison of the processes involved in DR, IR, image scoring and relatedness (or kin selection in the broad sense), and of the conditions in which they can account for cooperation. Arrows show donations. When an individual performs a donation (‘cooperates’), it pays a cost *c* and provides a benefit *b* to another individual. DR brings a direct fitness benefit when partners respond by cooperating and when the probability of interactions ending *w* is sufficiently small. IR is expected to have the corresponding condition, but where returns come from individuals other than the recipient of the actor’s donation. Image scoring (Nowak & Sigmund, 1998) is a postulated mechanism of IR which brings a direct fitness benefit when the probability of knowing the image score of a recipient (*q* or ‘social acquaintanceship’ (Rand & Nowak, 2013)) is sufficiently large (Nowak & Sigmund, 1998). Finally, where strategies help copies of the same strategy, they can be said to be related, and cooperating brings an indirect fitness benefit according to Hamilton’s rule. Note the similar form of these conditions.

IR has consequently become established as a “decisive” (Nowak & Sigmund, 1998) explanation for cooperation, and a “defining” (Yoeli et al., 2013) aspect of human sociality, accounting for large-scale fluid societies. In particular, IR is considered to be the reason why individuals care about being seen to help others, i.e. about their ‘reputations’, e.g (Brandt & Sigmund, 2005; Engelmann & Fischbacher, 2009; Fehr & Fischbacher, 2003; Fu et al., 2008; Ghang & Nowak, 2015; Milinski, 2016; Milinski et al., 2002a, 2002b). As well as being linked with the evolution of language (Nowak & Sigmund, 1998), it is further claimed that IR “constitute[s] the biological basis of morality (Pacheco et al., 2006); and that it “may have provided the selective challenge driving the cerebral expansion in human evolution” (Nowak & Sigmund, 2005). The literature on IR has therefore burgeoned (e.g.(Brandt & Sigmund, 2004, 2005; Hilbe et al., 2018; Ohtsuki & Iwasa, 2004, 2006; Pacheco et al., 2006; Panchanathan & Boyd, 2003, 2004; Santos et al., 2018; Suzuki & Akiyama, 2008; Takahashi & Mashima, 2006; Tanimoto, 2007; Uchida & Sigmund, 2010; Whitaker et al., 2016; Whitaker et al., 2018).

Here, I point out that classic models of IR are actually models of relatedness, or ‘kin selection’ in the broad sense of sharing genes more than average members of a population, whether by common ancestry or some other mechanism (West et al., 2007). Agents in these models do not help other helpers to receive something back from third parties; instead, they help because this furthers the spread of a strategy of helping those who help others. This can be understood in terms of Hamilton’s inclusive fitness theory: I propose that rather than getting direct fitness returns through indirect reciprocity, helpers are getting indirect fitness returns through their relatedness to recipients (Frank, 1998; Hamilton, 1964; Marshall, 2015; Queller, 2011). Unfortunately, authors have not always recognized the role of relatedness: a recent review of the most highly cited modelling papers on altruism found that all of them had wrongly ascribed their results to other factors such as spatial structure (Kay et al., 2020). In this paper, I show how authors have failed to identify the role of relatedness in models of indirect reciprocity.

### The problems with image scoring

Image scoring has long been known to be unstable with respect to ‘second order’ strategies like standing which consider how a recipient has previously behaved (Leimar & Hammerstein, 2001). However, this has been interpreted as meaning that image scoring is a weaker strategy of IR. My point in this paper is that image scoring is not a strategy of IR at all but is a strategy of helping relatives. To see this, we need to analyze how image scoring operates. For cooperation (as referred to in game theory as a behaviour which benefits others but which may involve a cost to the actor (West et al., 2007) to bring fitness benefits, there must be a relationship between giving and receiving so that the costs of cooperating are outweighed by the benefits of receiving, relative to non-cooperators. Image scoring offers a mechanism for linking donations with returns by identifying those who have cooperated and directing cooperation towards them. This is because image scorers are discriminating: they cooperate according to the recipient image score, which is a function of how cooperative the recipient itself has been. However, the logic of image scoring contains an in-built inconsistency which means it cannot explain IR.

This inconsistency centres on the fact that image scoring strategies do not evolve when individuals are self-interested, i.e. when they are maximizing their payoff. A population of image scorers is invaded by those who help others indiscriminately (Leimar & Hammerstein, 2001). This is because when a population is dominated by individuals playing a strategy of ‘help those with a positive image score’, an individual will receive more when they take more opportunities to cooperate. Thus, it will pay individuals to help on every move, even if this means helping defectors rather than playing the discriminating image scoring strategy. Hence, image scoring is not in an individuals’ best interests.

### Evidence for helping those who help others

Despite this theoretical result, it is widely thought that people do engage in image scoring. Yet empirical evidence supporting image scoring as an adaptive strategy may be weaker than often supposed. There is evidence that people behave more prosocially when they are observed, albeit in a more nuanced way than commonly thought (Bradley et al., 2018). However, this cannot be taken as evidence for IR as opposed to the range of other possible benefits arising from being observed, such as reputation-based partner choice (Roberts, 1998; Roberts et al., 2021; Sylwester & Roberts, 2010, 2013)). Students in experimental economic games do show at least partial discrimination, tending to give more often to those with high image scores (Wedekind & Milinski, 2000). However, while this has been interpreted as evidence for IR via image scoring, the experiments do not necessarily show that those who donate more get higher payoffs. Groups with more donors can do better (Milinski et al., 2002b; Wedekind & Milinski, 2000), but I point out that this has to be true because the benefit of receiving a donation is always set by the experimenter to be greater than the cost of donating (otherwise reciprocity would bring no advantage), hence I propose that the relationship between donations and group profit is an input rather than an interesting result of the experiment. IR requires that individuals do better than others within their own groups by engaging in donations, but there is little evidence for this. Thus, there appears to be no clear empirical evidence that image scoring is an adaptive strategy of IR.

### Is image scoring still relevant?

Image scoring is against individual strategic interests, leading to a conflict of interest between individual and strategy. Theorists have developed alternative strategies including ‘standing’ strategies (Leimar & Hammerstein, 2001; Sugden, 1986) and the ‘leading eight’ set of strategies (Ohtsuki & Iwasa, 2006). Such strategies take a more sophisticated view of defections so that agents do not lose their good standing upon ‘justified’ defection (Panchanathan & Boyd, 2003). So why does understanding image scoring remain important? First, the classic models of image scoring (Nowak & Sigmund, 1998) set in train a large theoretical and empirical field (Nowak & Sigmund, 2005) that was explicitly based on the interpretation of image scoring as being IR (Nowak & Sigmund, 2005). If we are to learn from these models of IR then we must understand whether they really are based on reputation at all. Secondly, despite many developments, the image scoring paper continues to be by far the most heavily cited paper on IR (e.g. (Henrich & Muthukrishna, 2021; Hilbe et al., 2018; Whitaker et al., 2018)). Thirdly, whilst image scoring models lack robustness due to the incentives involved, higher order strategies like standing have been criticized for their unrealistic informational requirements (Clark et al., 2020). Although a recent paper attempts to find a stable model of indirect reciprocity (Clark et al., 2020), this required modelling a payoff structure different to that of reciprocal help. Fourthly, empirical work on ‘helping those who help others’ remains much more relevant to image scoring than to more sophisticated strategies (e.g. (Milinski et al., 2001). Finally, I argue that the nature and significance of image scoring-like strategies is currently under-appreciated. This may seem paradoxical, but it is because I propose that image scoring can work, albeit as a strategy of helping relatives, not one of IR.

### Helping in image scoring systems works through relatedness not reciprocity

I propose that helping in image scoring systems is explained by relatedness. Individuals help other helpers, not because of reciprocal help, but because by doing so they help copies of the “help those who help others” strategy. Thus, it is a mistake to look for direct fitness benefits through reputation and indirect reciprocation. Instead, individuals get indirect fitness benefits which accrue when a helper is related to a recipient. The difference between strategies that depend on repeated interactions and those that depend on relatedness is visualized in Figure 3.

Inclusive fitness effects like this can be viewed from the perspective of the indirect fitness gains to the individual or from the perspective of the fitness gains to the strategy. Evolutionary simulations use the payoff accrued to strategies to determine how many individuals in the next generation carry that strategy: this is how strategies evolve and how we find which strategies are evolutionarily stable (Leimar & Hammerstein, 2001; Nowak & Sigmund, 1998; Roberts, 2008). The problem arises because individuals playing image scoring by helping those who help others are actually helping copies of their own strategy as carried by their recipients in which case the payoff to the helping strategy *increases* with every act of help (Figure 2). Image scoring can therefore be seen to be a form of relatedness in which the image scores are markers showing that individuals share the ‘help those who help others’ strategy

**Figure 2.**
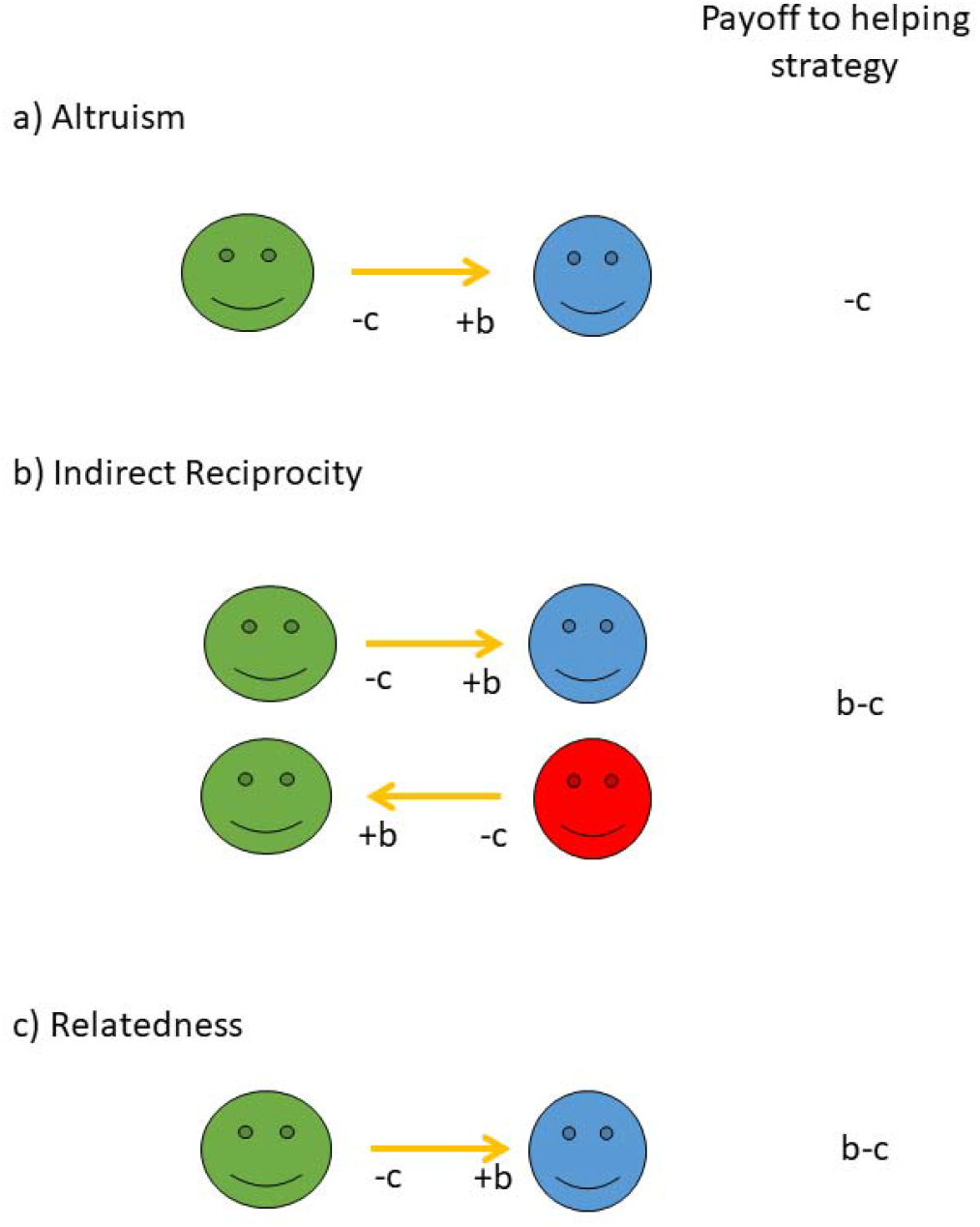
Calculating payoffs to helping strategies in simulations. In (a), altruism involves paying a cost and benefiting another individual. One way to explain this is through IR (b). Here, the individual makes a net benefit provided b>c and with the simplifying assumption that every act is reciprocated. The problem highlighted here is that relatedness (c) results in similar net benefits whenever the recipient shares the same strategy. This is because in computer simulations, the net payoffs to strategy members are summed and used to calculate the proportion of each strategy for the next generation. Thus, mechanisms (b) and (c) may be indistinguishable in typical simulations.

**Figure 3.**
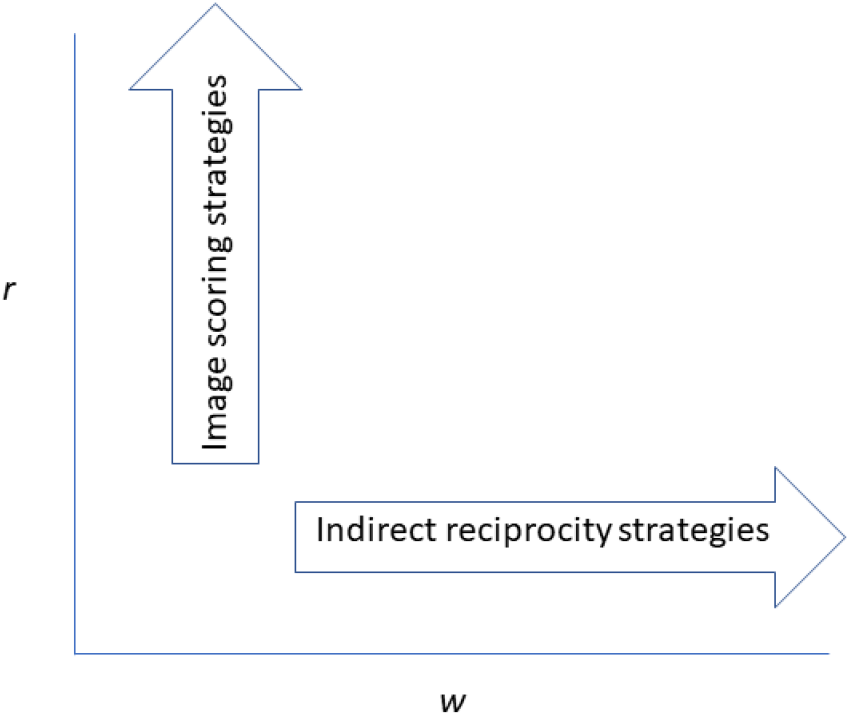
The roles of relatedness and of repeated interaction. The figure illustrates how image scoring strategies depend on relatedness *r* whilst indirect reciprocity strategies depend on the probability of repeat interactions (between third parties) *w*, and that these are separate dimensions.

If image scoring systems are actually driven by relatedness and not reciprocity then the condition for helping is:

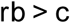

namely when the costs *c* to the actor are more than offset by the benefits *b* to a recipient scaled by their relatedness *r* (Hamilton, 1964). The commonest way in which altruism evolves is through genealogical relatedness. However, relatedness quantifies the genetic similarity between any two individuals (relative to the average similarity of all individuals in the population (West et al., 2007) and is not confined to cases of close genealogical descent. Hamilton’s rule takes the same form as the condition for cooperation in image scoring systems derived by Nowak and Sigmund, namely:

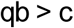

where q is ‘social acquaintanceship’ (or the probability of knowing the image score of a recipient) (Nowak, 2006; Rand & Nowak, 2013). The point I wish to make here is that this equation is not just similar to Hamilton’s rule, as remarked upon by the authors, but it is the same, because social acquaintanceship is equivalent to relatedness. Agents pay the cost of helping when this is offset by the benefit to the recipient multiplied by their chance of sharing the same strategy. I am suggesting that this is selected for through indirect fitness benefits, and not through IR. One corollary of this is that image scoring does not require repeated interactions and therefore is not a strategy of reciprocity.

The impact of relatedness can be seen in the simulations of (Leimar & Hammerstein, 2001). If helping behaviour is a function of relatedness rather than reciprocity, then donations should increase as average relatedness increases. One way to vary relatedness in models is to vary the demographic structure. (Leimar & Hammerstein, 2001) modelled IR using an island structure, ostensibly to reduce the effect of genetic drift. Whereas in a single population, helping another individual of the ‘help those who help others’ strategy means the benefit goes entirely to the reproductive success of the actor’s (and recipient’s) strategy, if the next generation is formed of only a portion of a local population, then the relatedness of a recipient will effectively be reduced. Thus, as populations become more divided into islands, so it becomes harder to satisfy the condition *rb > c* for helping. This can be seen in their simulations where island structure reduces image scoring strategies. In their paper, they attribute this effect to genetic drift but it can be better understood as an effect of declining relatedness in island populations. Notably they found that the standing strategy was unaffected by the population structure. I interpret this as being because the standing strategy does not require relatedness to work.

### Helping as a green beard

A corollary of my proposal that image scoring operates through relatedness is that image scores are not reputation indices but are actually kinship cues. Where individuals are not close kin, they may act as ‘green beards’: they display a distinguishing phenotype (helping others) allowing individuals to direct help to those who are genetically related (Dawkins, 1976; West & Gardner, 2010). Green beard altruism is expected to be rare (but see e.g. (Keller & Ross, 1998)), because individuals might ‘cheat’, displaying a green beard when they do not have the altruistic gene. However, as recognised by Dawkins himself, helping others is a uniquely honest green beard in that the expression of the helping behaviour is the green beard itself. So helping those who help others is a special case that can promote the spread of altruistic behaviour through the green beard effect of recognizing and favouring those that share the strategy, even when they are not genealogical kin.

It has long been recognized that if individuals help those displaying the green beard of helping, this becomes directly reciprocal help (Biernaskie et al., 2011; Dugatkin et al., 1994; Nee, 1989; Price, 2006; Trivers, 2006). In other words, strategies like like Tit-for-Tat in models of direct reciprocity (DR), are generally interpreted as involving the exchange of net direct benefits, but this may be difficult to distinguish from a green beard effect of helping those who help you. Nevertheless, most authors would consider that DR can be distinguished in real-world interactions (e.g. (Wilkinson, 1984) and the above papers interpreting DR in terms of green beards represent a minute proportion of the literature on DR.

None of these papers mention IR at all, but it might reasonably be thought that the same arguments about the interchangeability of direct and indirect fitness effects should apply to both types of reciprocity. I point out here that there is a fundamental difference between the two (Figure 4). In DR, if you help another DR agent then you are assured of getting *both* the direct fitness benefits of reciprocation *and* the indirect fitness benefits of helping another DR strategist (keeping in mind that we should avoid double counting and that we should strictly speaking be specific about the interests of the genes versus the individuals). However, in the IR scenario, if you help another IR agent then there can be a separation of direct and indirect benefits so you will get the indirect benefits if the recipient is an IR, but you may not get a return from another individual unless another image scorer observed the donation. This means, crucially, that following an image scoring strategy can result in indirect benefits but not direct ones and therefore that image scoring may work as a strategy of helping relatives without being a stable strategy of reciprocity.

**Figure 4.**
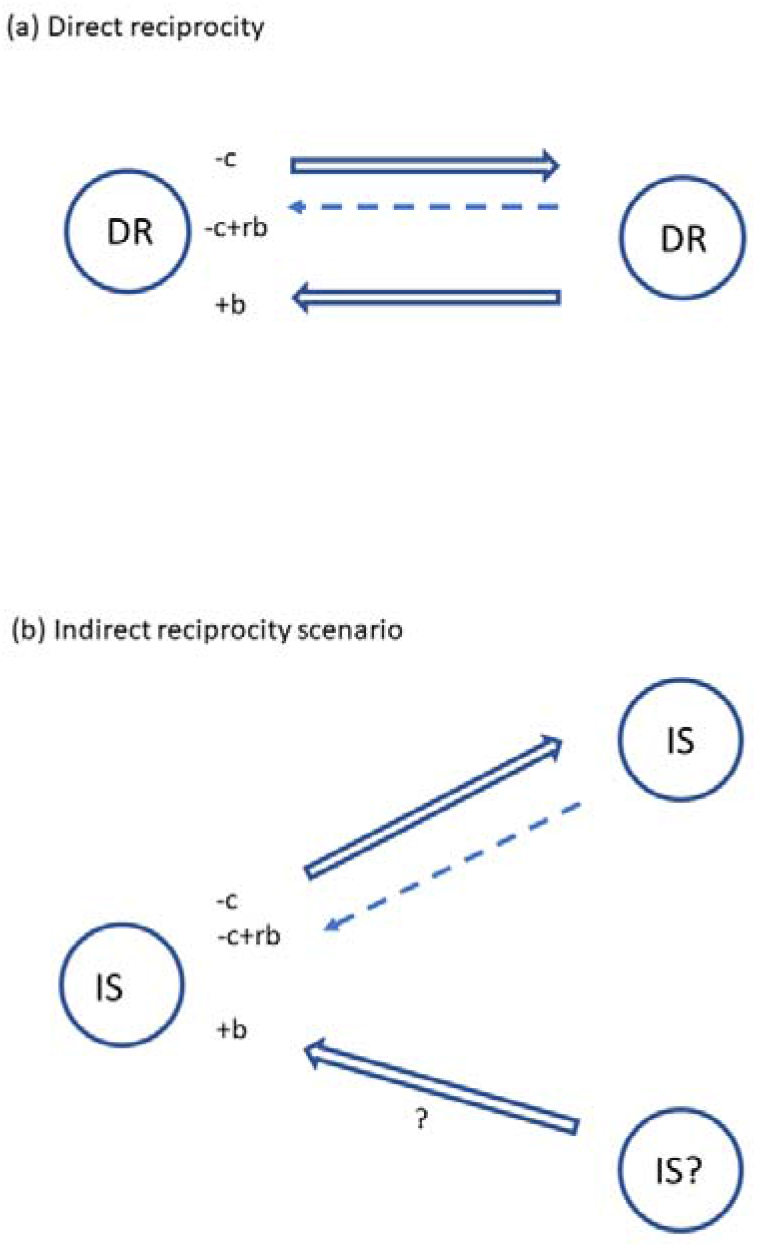
Direct and indirect fitness effects in direct reciprocity and image scoring. Consider the payoffs to the left hand individual. In (a) a direct reciprocity agent (DR) pays a cost *c* to help (full arrow) and receives a benefit *b* from another DR agent. This gives the same direct fitness payoff as if we consider that the left hand agent gets an indirect fitness benefit from helping another agent sharing the same strategy (relatedness *r* = 1). However, in (b) there is a separation between the individual that is helped and the chance of another individual reciprocating. This means that an image scoring (IS) agent helping another IS agent is assured of the indirect fitness benefits but may or may not make a direct fitness return.

### Are higher order strategies of indirect reciprocity also reliant on relatedness?

What about the more sophisticated strategies like standing (Leimar & Hammerstein, 2001; Sugden, 1986), stern judging (Pacheco et al., 2006) and others of the ‘leading eight’ (Ohtsuki & Iwasa, 2006)? Such strategies can be evolutionarily stable and can in theory allow IR by bringing direct fitness benefits. However, just as with DR, they could be interpreted in terms of indirect fitness benefits rather than direct ones when these are equivalent. Hence it is possible that those following a stern judging strategy, for example, may do well because they are good at directing help at other carriers of the same strategy. Indirect benefits could enable these strategies to become established below the threshold of repeated interactions among third parties where they would become stable IR strategies. If indeed indirect fitness benefits are important then strategies like stern judging or the ‘social comparison heuristic’ (Whitaker et al., 2016) may be less to do with morality, reputations and norms (as presented in (Santos et al., 2018) and more to do with being good at directing help to relatives.

## Conclusion

To summarize, what has been termed ‘image scoring’ in a large literature (Henrich & Muthukrishna, 2021; Okada, 2020) cannot be explained by individual adaptation; does not explain why help is directed discriminatingly; is not a form of reciprocity; and is actually a function of relatedness. It works not because of getting a reciprocal benefit but by helping copies of the strategy of “help those who help others”. Recognizing this resolves the paradox of how cooperation in image scoring systems is individually costly yet beneficial to the strategy: donating is only beneficial in inclusive fitness terms. Reciprocity may be an illusion resulting from individuals receiving after revealing their strategy by helping.

IR has become synonymous with a concern for reputation e.g. (Henrich & Muthukrishna, 2021). If helping those who help others is driven by relatedness rather than reputation, then this focus may be misplaced. This is very important because so much store is set by authors claiming that reputation is a key part of what makes us human (e.g. ((Henrich & Muthukrishna, 2021; Nowak & Sigmund, 2005; Rand & Nowak, 2013; Whitaker et al., 2018; Yoeli et al., 2013)). There is nevertheless evidence that people are more helpful when observed. Another theory for how building a cooperative reputation might be advantageous that has received less attention is reputation-based partner choice. This differs from IR in that cooperative reputations bring benefits through access to profitable partnerships (Roberts, 1998; Roberts et al., 2021; Sylwester & Roberts, 2010). Theory and evidence suggest that reputations can be a powerful mechanism supporting cooperation but that IR plays little if any role (Sylwester & Roberts, 2013).

The role of inclusive fitness in sociality has recently been challenged ((Nowak et al., 2010) but see also ripostes from e.g. (Abbot et al., 2011)), yet I have argued here that image scoring systems actually work through kin selection. Given that image scoring has been cited as a decisive factor in the evolution of human large-scale sociality including the evolution of language and of morality (Nowak & Sigmund, 2005; Santos et al., 2018) these issues must be fundamentally revisited. Strategies of helping those who help others may be facilitated by cultural evolution (Boyd & Richerson, 1989) because population structure pertinent to cultural transmission may be more favourable than that relevant to transmission of genes. Prosocial strategies of helping those who help others might therefore spread culturally. This possibility should be the focus of future work.

## Acknowledgements

I thank T. Sherratt, R. Boyd and J. Kendal for comments on earlier versions.

## Ethics approval

No ethical approval was required for this work.

## Consent to participate

n/a

## Consent for publication

n/a

## Availability of data and material

There are no associated data or materials.

## Code availability

n/a

## Authors’ contributions

GR was solely responsible for writing the manuscript.

